# A Novel Efficient Algorithm for Common Variants Genotyping from Low-Coverage Sequencing Data

**DOI:** 10.1101/2024.12.01.626280

**Authors:** Shixuan Zhang, Kanglong Xiao, Xinyi He, Meng Hao, Yanyun Ma, Yuxin Tian, Li Jin, Yi Li, Jiucun Wang, Yi Wang

## Abstract

Low-coverage whole-genome sequencing (LC-WGS) combined with imputation represents a cost-effective genotyping strategy for genome-wide association studies (GWAS) in population genetics. In this study, the Limpute algorithm was developed specifically for genotyping from low-coverage sequencing data, it extracts variant information from low-coverage sequencing data by the novel virtual probes and subsequently performs imputation through cross-reference between samples. Compared to the currently dominant algorithm for low-coverage sequencing data, GLIMPSE2, Limpute achieved similar imputation performance within common variants (r^2^>0.87) while the GLIMPSE2 has a runtime approximately five times longer than that of the Limpute. Furthermore, to fully evaluate the accuracy of genotype imputation by Limpute, we utilized high-coverage whole-genome sequencing data (30x), microarray data, and high-coverage whole-exome sequencing data (30x) as validation sets respectively. The results demonstrated that Limpute has a good imputation performance for common variants using low-coverage sequencing data (1x: r^2^ > 0.87; 3x: r^2^ > 0.92; 5x: r^2^ > 0.93). In summary, we present a highly efficient, low-cost algorithm for genotyping from low-coverage sequencing data, offering substantial support for genetic research.

## Introduction

Genome-wide association analysis studies (GWAS) provide opportunities to explore the genetic susceptibility of diseases (1-5), more and more research highlighting the advantages of low-coverage whole-genome sequencing (WGS) (4-8). High-coverage sequencing in large sample sizes incurs high costs, leading to the proposition of low-coverage WGS as an economically efficient alternative (11, 12). Studies have shown that systematic low-coverage sequencing can capture a similar number of common variants as standard SNP array platforms and a greater number of rare variants (13-15), complementing genetic signals missed by SNP array platforms (14, 16).

While low-depth sequencing offers a higher cost-effectiveness (17), the probabilistic nature of the data and high missing rates challenge the credibility of current GWAS studies (18). Imputation algorithms utilize information from haplotype reference panels to fill in sparse mapping read gaps (19, 20). In 2023, Simone Rubinacci et al. introduced the GLIMPSE2 algorithm, which significantly enhances the imputation performance of variant sites based on low-coverage sequencing data with Biobank reference panels (14, 21). However, there are still drawbacks regarding high computational costs and time consumption. Given the demands of genetic research in large-scale population cohorts for GWAS studies, genotype imputation algorithms need to maintain high performance while minimizing computational resource consumption as much as possible.

In this study, a novel genotype likelihood imputation algorithm, Limpute, was developed and its genotyping performance was evaluated (supplementary fig. S1). When microarray data were used as the validation set, the genotyping performance of Limpute within common variants (r^2^>0.87) was comparable to the GLIMPSE2(21) algorithm, but the runtime was reduced by approximately 88%. Furthermore, Limpute outperformed the Beagle v.5.4(22) algorithm in both runtime and genotyping performance. Additionally, high-coverage whole-genome sequencing data (30x), microarray data, and high-coverage whole-exome sequencing data (30x) were used as validation sets respectively to fully evaluate the accuracy of genotype imputation by Limpute. The results indicated that the squared pearson correlation coefficients (r^2^) of the Limpute algorithm for low-coverage sequencing data (1x, 3x, 5x) exceeded 0.80 (1x: r^2^ > 0.87; 3x: r^2^ > 0.92; 5x: r^2^ > 0.93), demonstrating a high level of precision. Overall, this study suggests that the Limpute algorithm is a viable method for genotyping with low-depth sequencing data.

## Results

### 1. Limpute: A Novel Framework for Imputation in Low-Coverage Sequencing Data

The overview of the Limpute algorithm is as illustrated (Fig. 1). Initially, the input to Limpute is a genotype likelihood matrix, which is derived from low-coverage sequencing data by the virtual probe, termed GoldProbe. Subsequently, the obtained sample genotype likelihood matrix is partitioned into two segments: one serving as the reference and the other as the target. Employing a Hidden Markov Model (HMM) (23), the Limpute algorithm facilitates cross-reference between samples, thereby outputting a genotype probability matrix for the target samples. This matrix can then be converted into a VCF file for the target samples. Notably, the designation of reference and target samples is customizable based on the specific application. The efficiency of this approach stems from two pivotal algorithmic features: Firstly, the bit-vector algorithm of the virtual probe is leveraged to extract information correlated to corresponding loci from the sample sequencing data, which is then stored using a hash index structure. This approach ensures accurate information extraction while enhancing computational speed. Secondly, by utilizing the genotype likelihood matrix for cross-reference between samples, the genotype calling is directly accomplished, bypassing the need to generate memory-intensive genome alignment files (BAM files). It not only accelerates the process but also reduces memory consumption.

**Fig. 1.**
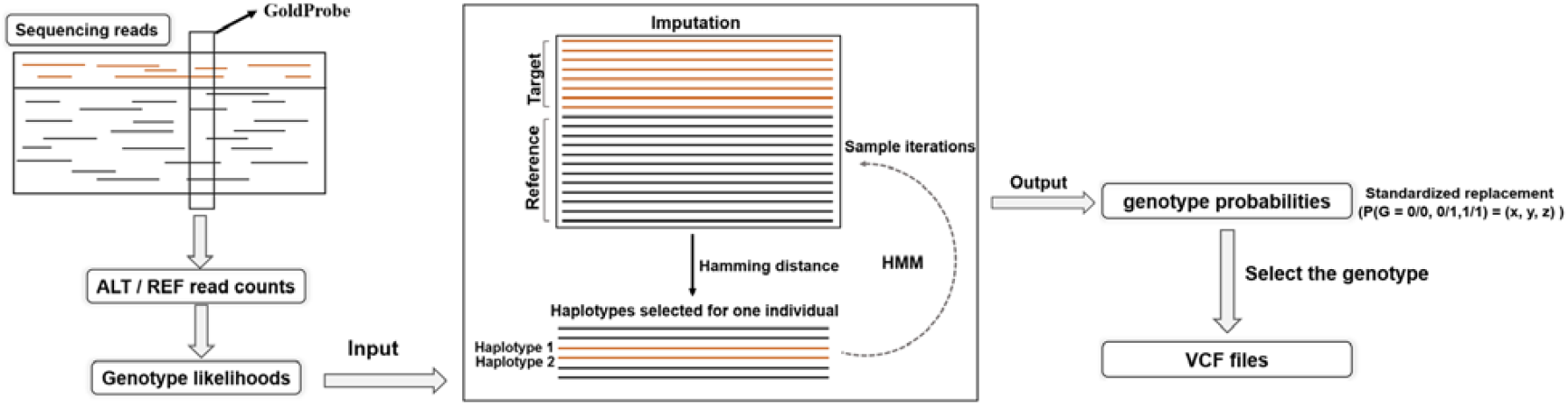
The overview of the Limpute algorithm. The input to this algorithm comprises the counts of reference and variant reads at all SNP loci, obtained from sequencing data by utilizing the virtual probe, GoldProbe, which subsequently facilitates the generation of a genotype likelihood matrix. The Limpute algorithm then employs a cross-reference between samples, utilizing the Hamming distance and the Hidden Markov Model (HMM) to refine the genotype likelihoods. This iterative process culminates in the output of genotype probabilities for the target samples, producing corresponding sample variant call format (VCF) files.

### 2. Comparison of Imputation Performance Between Three Algorithms

Two widely recognized genotyping algorithms are GLIMPSE2 and Beagle v.5.4. We compared the imputation performance of Limpute against these algorithms using genotype accuracy and squared pearson correlation coefficients (r^2^). Using microarray data as the validation set, we evaluated 323,649 SNP loci on chromosome one. The results indicated that the genotype accuracy of the Limpute algorithm (1x: 0.967; 3x: 0.976; 5x: 0.977) was similar to that of the GLIMPSE2 algorithm (1x: 0.969; 3x: 0.975; 5x: 0.976) (Fig.2 A, B, C; supplementary table S1). Moreover, the squared pearson correlation coefficients of the Limpute algorithm (1x: r^2^ > 0.87; 3x: r^2^ > 0.90; 5x: r^2^ > 0.91) at common variant SNPs (MAF > 5%) were comparable to that of the GLIMPSE2 algorithm (1x: r^2^ > 0.88; 3x: r^2^ > 0.89; 5x: r^2^ > 0.90) (Fig2. D, E, F; supplementary table S1). Meanwhile, both the Limpute and GLIMPSE2 algorithms demonstrated superior imputation performance compared to the Beagle v.5.4 algorithm (1x: 0.816; 3x: 0.913; 5x: 0.948).

**Fig. 2.**
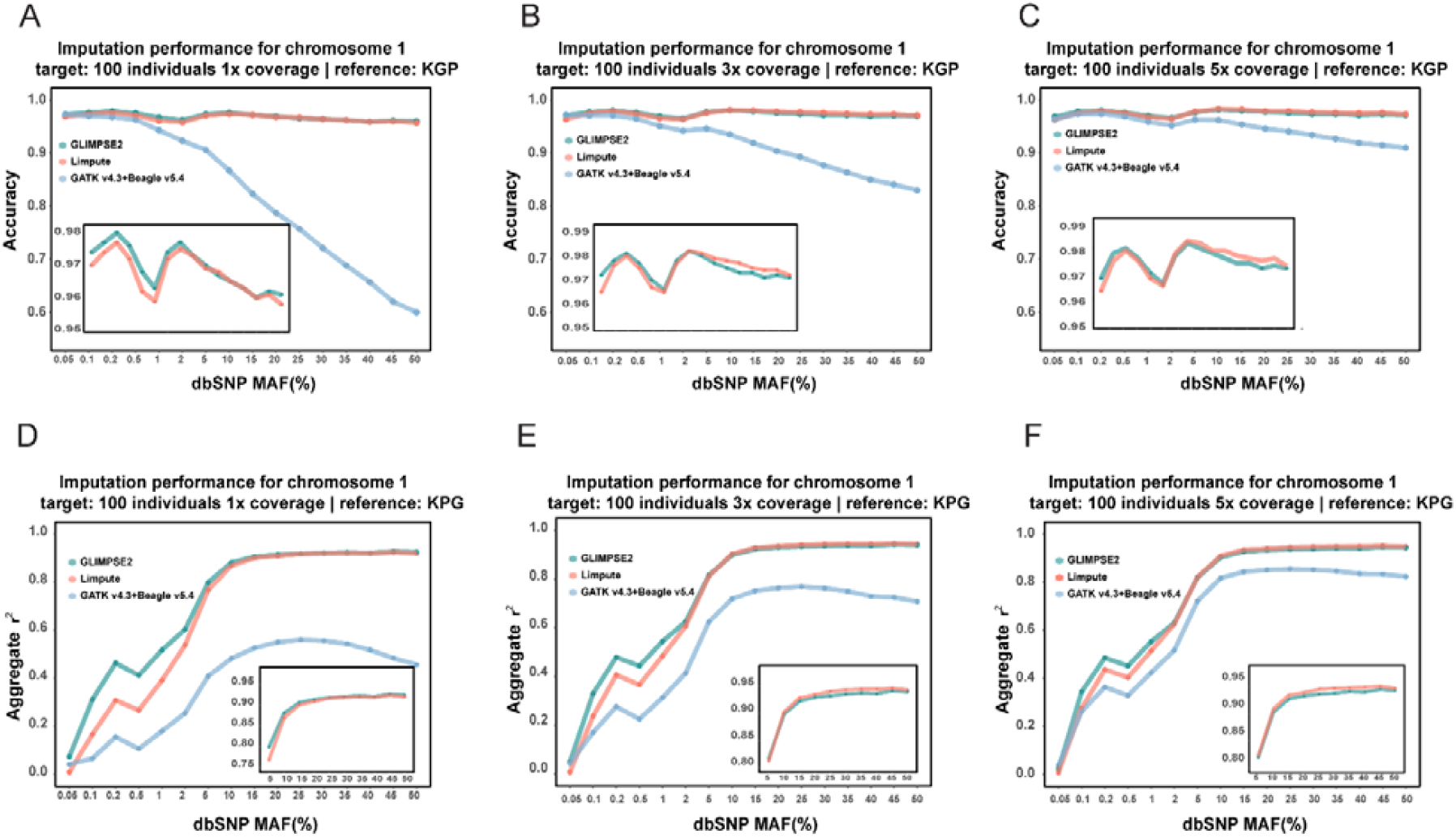
Performance measurement of three algorithms for genotype imputation. **A:** Line graphs showing the distribution of genotype accuracy for the Limpute algorithm, Beagle v.5.4 algorithm, and GLIMPSE2 algorithm using 1x sequencing data. **B:** Line graphs showing the distribution of genotype accuracy for the Limpute algorithm, Beagle v.5.4 algorithm, and GLIMPSE2 algorithm using 3x sequencing data. **C:** Line graphs showing the distribution of genotype accuracy for the Limpute algorithm, Beagle v.5.4 algorithm, and GLIMPSE2 algorithm using 5x sequencing data. **D:** Line graphs showing the distribution of squared Pearson correlation coefficients (r^2^) for the Limpute algorithm, Beagle v.5.4 algorithm, and GLIMPSE2 algorithm using 1x sequencing data. **E:** Line graphs showing the distribution of squared Pearson correlation coefficients (r^2^) for the Limpute algorithm, Beagle v.5.4 algorithm, and GLIMPSE2 algorithm using 3x sequencing data. **F:** Line graphs showing the distribution of squared Pearson correlation coefficients (r^2^) for the Limpute algorithm, Beagle v.5.4 algorithm, and GLIMPSE2 algorithm using 5x sequencing data. **Notes:** The horizontal coordinates in each graph indicate the MAF interval segments (assessed SNP loci are divided according to the MAF values from the public database dbSNPs), and the vertical coordinates indicate the algorithm imputation performance. The inset figure provides a clearer comparison between the Limpute algorithm and the GLIMPSE2 algorithm.

### 3. Comparison of Memory Consumption and Runtime Between Three Algorithms

The comparison of memory consumption and runtime across three genotype imputation algorithms indicates that the Limpute algorithm exhibits a clear advantage in both aspects when obtaining genotypes at identical SNP loci (Fig. 3A, B). On the one hand, GLIMPSE2 and GATK v.4.3 plus Beagle v.5.4 consume approximately the same amount of physical memory (Fig. 3A), while the Limpute algorithm reduces memory usage by 66.85% compared to these algorithms. On the other hand, the overall runtime of the Limpute algorithm is reduced by approximately 87.93% compared to GLIMPSE2 and by approximately 91.13% compared to GATK v.4.3 plus Beagle v.5.4 (Fig. 3B). Overall, the Limpute algorithm demonstrates superiority in both memory consumption and runtime.

**Fig. 3.**
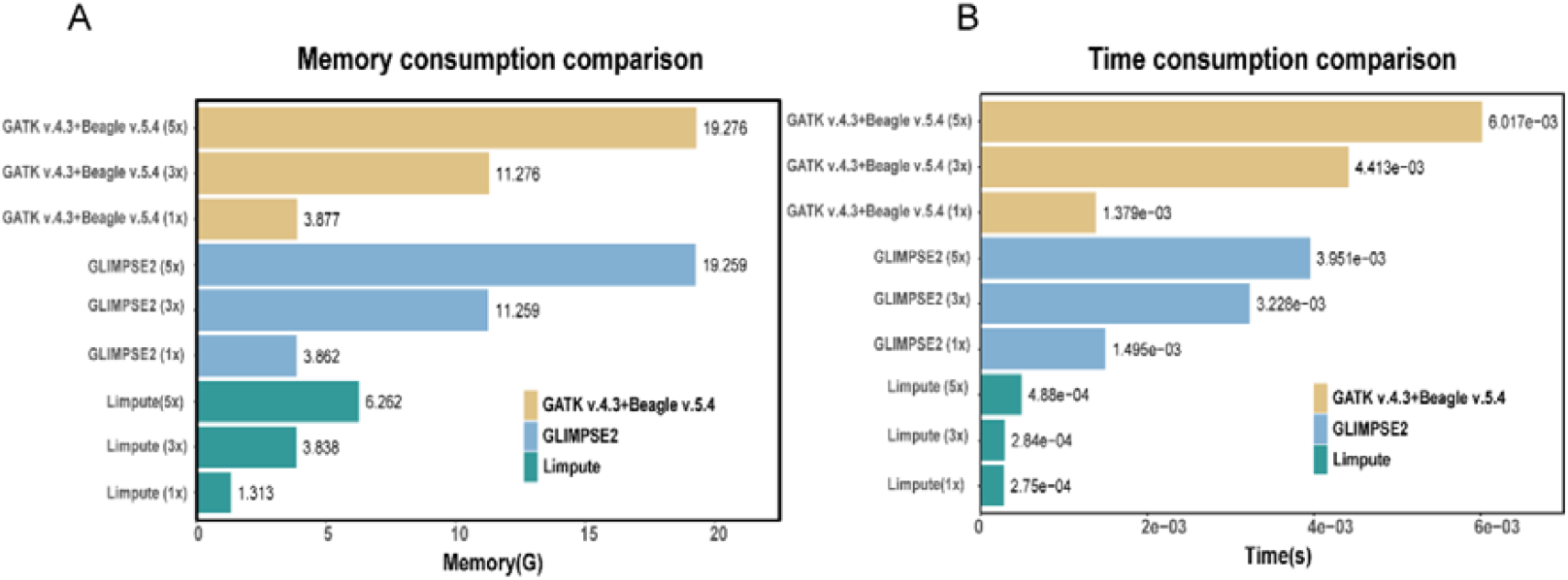
Comparison of memory consumption (A) and runtime (B) among the three algorithms. **A:** Comparison of the memory usage associated with intermediate files generated by the three algorithms during genotype calling across all SNP loci for a single sample. **B:** Comparison of the time required by the three algorithms to perform genotype calling for a single SNP locus in an individual sample.

### 4. Good Imputation Performance Within the Whole Exon Regions

Based on 671 Chinese subjects with high-coverage (30x) whole-exome sequencing data as the validation set, we evaluated the imputation accuracy of the Limpute algorithm for low-coverage (5x) whole-genome sequencing data, including 90,421 SNP loci. The results indicate that the Limpute algorithm achieved an average accuracy of 0.93 across all SNPs, with a squared pearson correlation coefficient reaching 0.83. Furthermore, we delved into the genotyping performance across different minor allele frequency (MAF) bins, revealing that the Limpute algorithm maintained an accuracy greater than 0.90 in each MAF segment (Fig.4A; supplementary table S2). Specifically, for common variant SNPs (MAF > 5%), the squared pearson correlation coefficient surpassed 0.83 (Fig.4B; supplementary table S2). In summary, the Limpute algorithm exhibits robust imputation performance within whole-exome SNP loci.

**Fig. 4.**
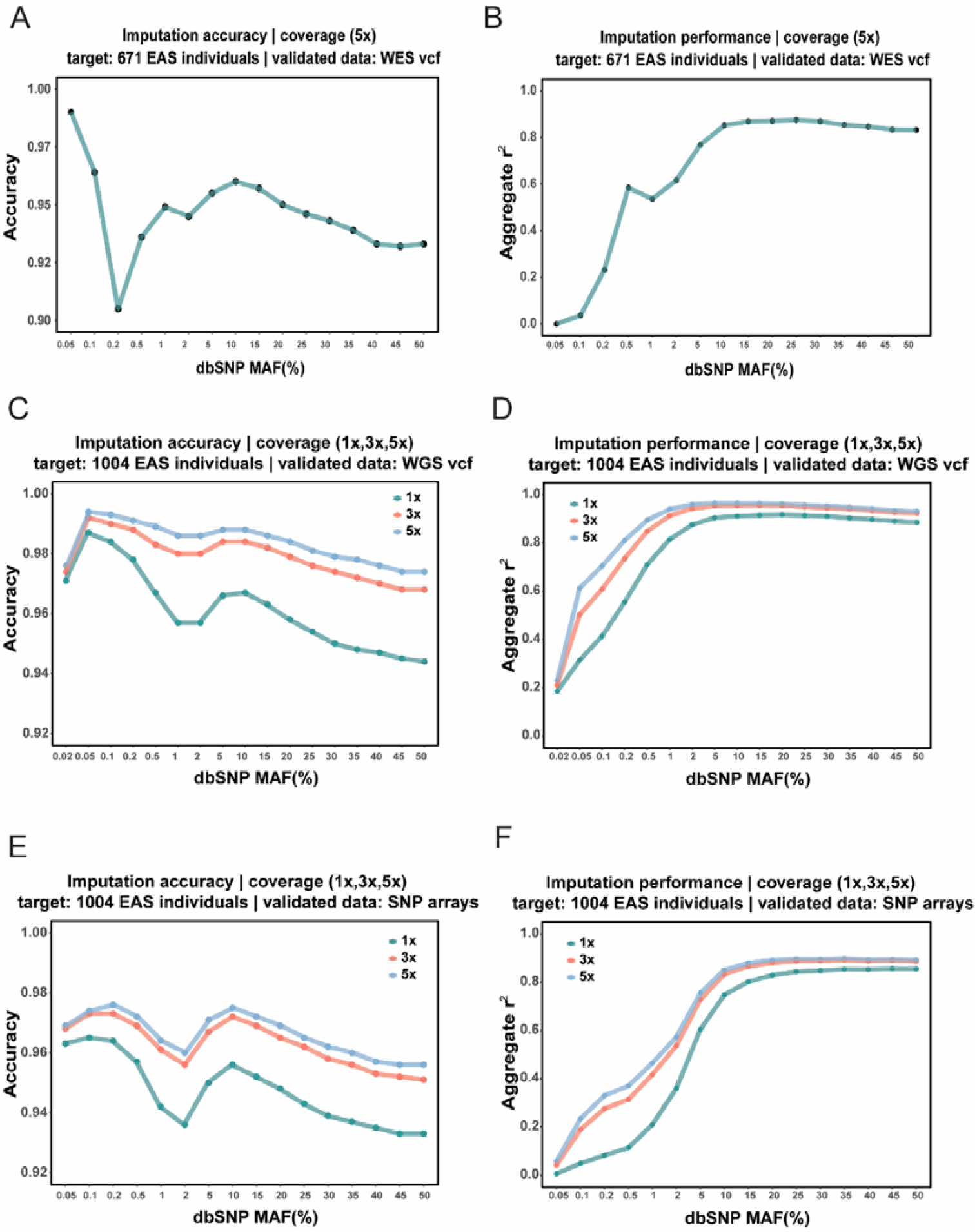
Genotyping performance measurement of Limpute algorithm. **A:** the line graph showing the genotyping accuracy of SNPs using low-coverage (5x) whole-genome sequencing data with the Limpute algorithm, validated against whole-exome data. **B:** the line graph showing the squared Pearson correlation coefficients of SNPs using low-coverage (5x) whole-genome sequencing data with the Limpute algorithm, validated against whole-exome data. **C:** the line graph showing the genotyping accuracy of SNPs using low-coverage (1x, 3x, 5x) whole-exome sequencing data with the Limpute algorithm, validated against whole-genome data. **D:** the line graph showing the squared Pearson correlation coefficients of SNPs using low-coverage (1x, 3x, 5x) whole-exome sequencing data with the Limpute algorithm, validated against whole-genome data. **E:** the line graph showing the genotyping accuracy of SNPs using low-coverage (1x, 3x, 5x) whole-exome sequencing data with the Limpute algorithm, validated against microarray data. **F:** the line graph showing the squared Pearson correlation coefficients of SNPs using low-coverage (1x, 3x, 5x) whole-exome sequencing data with the Limpute algorithm, validated against microarray data. **Notes:** The horizontal coordinates in each graph indicate the MAF interval segments (assessed SNP loci are divided according to the MAF values from the public database dbSNPs), and the vertical coordinates indicate the algorithm imputation performance.

### 5. Good Imputation Performance Within the Whole-Genome Regions

Based on high-coverage whole-genome sequencing data (30x) and microarray data from 1,004 Chinese volunteers, the accuracy of the Limpute algorithm for genotype calling of low-coverage exome sequencing data (1x, 3x, 5x) was evaluated. The results demonstrate that the Limpute algorithm performs well in genotype calling, consistent with the assessment of imputation performance in the exonic regions. Firstly, using high-coverage whole-genome sequencing data (30x) as the validation set, the imputation performance of 7,602,829 SNP loci were assessed. It shows that the accuracy of the Limpute algorithm exceeded 0.96 (1x: 0.964; 3x: 0.981; 5x: 0.986), with squared pearson correlation coefficients for common variant SNPs (MAF > 5%) all exceeding 0.88 (1x: r^2^ > 0.88; 3x: r^2^ > 0.92; 5x: r^2^ > 0.93) (Fig.4E; supplementary table S3). Secondly, using microarray data as the validation set, the imputation performance of 5,624,383 SNP loci were evaluated. It shows that the Limpute algorithm achieved an accuracy greater than 0.94 (1x: 0.946; 3x: 0.964; 5x: 0.967), with squared pearson correlation coefficients for common variant SNPs (MAF > 5%) all exceeding 0.72 (1x: r^2^ > 0.72; 3x: r^2^ > 0.81; 5x: r^2^ > 0.82) (Fig.4F; supplementary table S4). The genotype calling accuracy of the Limpute algorithm for SNPs segmented by different MAFs was consistently above 0.92 (Fig.4 C and E), and its performance improves with increasing sequencing depth.

## Discussion

This study presents an efficient genotype imputation algorithm for low-coverage sequencing, which performs genotype imputation by using cross-reference between samples. The imputation performance of the Limpute algorithm was evaluated on exome-wide SNP loci (671 samples) and genome-wide SNP loci (1004 samples), demonstrating a high level of genotype accuracy (r^2^>0.8). In another aspect, a comparison was made between the Limpute algorithm and two internationally recognized algorithms, Beagle v.5.4 and GLIMPSE2, in terms of genotype accuracy, runtime, and memory consumption. The results indicate that the genotype imputation performance of the Limpute algorithm within common variants (r^2^>0.87) is comparable to that of GLIMPSE2, and both significantly outperform the Beagle v.5.4 algorithm (Fig. 2; supplementary Table S1). Notably, due to the unique genotype imputation approach of Limpute, it achieves the best performance in terms of runtime and memory consumption among the three algorithms. Specifically, the Limpute algorithm reduces memory usage by 66.85%, decreases runtime by approximately 87.93% compared to GLIMPSE2, and by 91.13% compared to GATK v.4.3 plus Beagle v.5.4 (Fig. 3B), significantly lowering operational costs.

To achieve genotype imputation for low-coverage sequencing data, previous algorithms have primarily based on high-coverage reference panels, such as the 1000 Genomes Project (24) and gnomAD (25). GLIMPSE2 is a dominance algorithm of this type of algorithm, it has shown that the performance of GLIMPSE2 outperforms that of the GLIMPSE and the QUILT(21). In the present study, GLIMPSE2 demonstrated a good imputation accuracy (r^2^>0.88) for low-coverage sequencing data (Fig.2 A, B, C; Supplementary Table S2). However, The Limpute’s imputation performance is comparable to that of the GLIMPSE2 and drastically decreases operational costs, it reduces memory usage by 66.85%, decreases runtime by approximately 87.93% compared to GLIMPSE2.

This study proposes Limpute, a low-coverage sequencing genotyping algorithm with high imputation performance and low consumption. Limpute introduces innovative use of virtual probes during genotype imputation, eliminating the need to generate genome alignment files (27), which significantly reduces both physical memory usage and runtime. Its unique approach that cross-reference between samples, enables highly accurate genotype imputation. The algorithm’s imputation performance for low-coverage sequencing data within common variants is comparable to that of the GLIMPSE2 (21) algorithm and significantly outperforms Beagle v.5.4 (21). Additionally, Limpute achieves the best runtime and memory efficiency among the three algorithms. By ensuring high-accuracy genotype imputation while drastically reducing runtime and memory consumption, Limpute is well-suited to minimize research resource demands in large-scale population cohort genomic studies.

There are some limitations in our study. First, the genotype imputation performance of the Limpute algorithm for rare variant SNPs (MAF < 1%) is inferior to that of GLIMPSE2. Second, the sample size used in this study was relatively small, which may have limited the ability to fully demonstrate the advantages of Limpute’s unique genotype imputation approach. Lastly, the sample data utilized in this study were exclusively from East Asian populations, so the accuracy of genotype imputation in other populations remains to be validated. As low-coverage sequencing is likely to become more prevalent in future genomic research (2, 21), we offer a highly accurate, low-cost genotype imputation algorithm suitable for a wide range of genetic applications.

## Conclusion

In this study, the Limpute algorithm was developed specifically for genotyping from low-coverage sequencing data, it extracts variant information from low-coverage sequencing data by the novel virtual probes and subsequently performs imputation through cross-reference between samples. it ensures high-accuracy genotype imputation while drastically reducing runtime and memory consumption, Limpute is well-suited to minimize research resource demands in large-scale population cohort genomic studies.

## Materials and Methods

### Algorithm principle

#### a: The GoldArray Pipeline

The GoldArray is an innovative bioinformatics pipeline that mimics the conventional microarray system in the era of NGS. The GoldArray consists of millions of paired GoldProbes. Each probe in the pair is designed to detect the in silico hybridization signal for a SNP allele from raw sequencing reads. (We view sequencing reads as “DNA”).

A GoldProbe is a 63 bp virtual oligo centered at a SNP. An in silico hybridization is performed between the GoldProbes and a NGS read sequence in the sense of edit distance using Myers’ bit vector algorithm. An edit distance within two is considered as a hit.

The two GoldProbe (A and B) in a pair are designed to be competitive. If a read results an edit distance of 2 with A and an edit distance of 1 with B, it will only be counted as B’s signal. If a draw appears, neither A nor B counts signals.

To accelerate the in silico hybridization of millions of GoldProbes, we use an exact hash index structure. We split the 63 bp GoldProbe into three parts, each of which is 21 bp. Since both edit distance of two GoldProbe is allowed, according to the pigeonhole principle, for a hit at least one exact match of the three parts should be observed. We perform Myers’s algorithm only if the reads show an exact match with any 21 bp part with a GoldProbe.

The final report of GoldArray is the two-read count of each SNP, one for the reference allele and the other for the alternative allele. The GoldArray is fast due to the index structure and accurate due to Myers’s algorithm and competitive counting.

#### b: The Imputation Algorithm

Our imputation algorithm is based on Wang’s idea (23) which models a haplotype as the recombined offspring of two ancestral haplotypes. Instead of many ancestral parent modeling, such restricted and simplified model results in a much faster handcrafted HMM algorithm. As an improvement of this idea (23), a Hamming distance based proposal routine is employed. Given current parental haplotype A and B, we want to suggest a candidate haplotype C from the population to replace A. Our heuristic is to define a distance:

Distance(A→C,B)=2*Hamming(A,C)-Hamming(B,C).

The definition suggests that the new candidate should be similar to the current candidate and should be different from the other haplotype. The pseudo code is as follows:

Input a population of genotype likelihood. Randomized the initial guessed haplotypes and parents. While likelihood increases {

for each sample, search its four parental haplotypes{

for given iterations {

Propose a replacement of one haplotype using the aforementioned proposal routine. Calculate the proposal’s likelihood using genotype likelihood and HMM model.

Accept or reject the proposal using a MH sample.

}

Sample the offspring’s haplotype using Viterbi decoding.

}

accumulate sample genotype probabilities into real valued accumulators

}

#### c: report the final genotype probabilities into a VCF-like format

The Imputation Algorithm will produce genotype probabilities files(P(G = 0/0, 0/1,1/1) = (x, y, z)). IF x≠y≠z, then we will select the genotype corresponding to the position with the highest probability for normalization (e.g., x>y & x> z, select ‘0/0’); while in other conditions, if x=y≠z OR x≠y=z, we will take three steps to produce the final genotype. First, capturing pattern SNPs with the same probability of two genotypes(e.g., P(G = 0/0,0/1,1/1) = (0.1,0.4,0.4)); Second, use the P(G = 0/0, 0/1,1/1) distribution to create a Genotype-matrix of 100,000(Including 0/0, 0/1,1/1); Third, use the Python ‘random’ module to randomly select a value in Genotype-matrix to replace the current phenotype probability P(G = 0/0, 0/1,1/1).

### Data used

In this study, we utilized sequencing data from a total of 1675 Chinese volunteer samples (generated on the Illumina sequencing platform), comprising 671 samples with high-coverage whole-exome sequencing data (30x) and 1004 samples with both high-coverage whole-genome sequencing data (30x) and microarray data (https://www.cncb.ac.cn/). Low-coverage sequencing data were obtained by randomly sampling reads from the high-coverage datasets. For the evaluation of SNPs in both the whole-exome and whole-genome regions, the Limpute algorithm employed cross-reference between samples to derive sample genotype files. In the algorithm comparison section, for the sake of standardization, the Limpute algorithm, GLIMPSE2(21) algorithm, and Beagle v.5.4(22) algorithm utilized the Phase3 Chromosome 1 VCF file (24) from the 1000 Genomes Project as a reference panel for sample genotype calling, facilitating direct comparisons.

The generation process of the validation set VCF files is as follows. Firstly, high-quality sequencing data are aligned to the human hg19 reference genome using the BWA (Burrows Wheeler Aligner) software (26), resulting in alignment files (bam files). Following the removal of duplicates and recalibration of base quality scores, the GATK v.4.3 (27) command ‘haplotypecaller’ (gatk.broadinstitute.org) is employed with default parameters to obtain the whole-exome and whole-genome VCF files, which serve as the validation sets. We use the Picard tools (biotools: Picard tools) to convert diverse genomic coordinates.

### Genotyping Performance Assessment

We employed two methodologies to evaluate the genotyping performance of our algorithm. Firstly, we utilized the genotype accuracy method (28) to assess the proportion of concordance between the genotypes inferred by the algorithm and those in the validation set for identical SNPs across all samples:

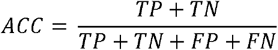

TP is the number of correct positive predictions, FP is the number of incorrect negative predictions, TN is the number of negative predictions that are correct, and FN is the number of positive predictions that are incorrect.

Secondly, the squared Pearson correlation coefficient (29) was utilized to compare the correlation between the genetic effect dosages inferred by the algorithm and those in the validation set, leveraging the tool aggRSquare (https://github.com/yukt/aggRSquare.git). Recognizing that the algorithm’s inference performance varies across SNPs with different minor allele frequencies (MAFs) (14), we utilized the MAF information of SNPs from the public database dbSNP to categorize these SNPs into 17 intervals based on their MAF values: (0–0.0002], (0.0002–0.0005], (0.0005–0.001], (0.001–0.002], (0.002–0.005], (0.005–0.01], (0.01–0.02], (0.02–0.05], (0.05–0.1], (0.1–0.15], (0.15–0.2], (0.2–0.25], (0.25–0.3], (0.3–0.35], (0.35–0.4], (0.4–0.45], (0.45–0.5]. Unless otherwise specified, all comparisons in this study pertained to SNPs located on the 22 autosomes.

### Time-Memory Consumption Evaluation Method

In the process of generating a genotype calling file for a sample from sequencing data, we individually tallied the total physical memory consumed by the Limpute algorithm, GLIMPSE2 algorithm, and the combination of GATK v.4.3 and Beagle v.5.4 algorithm in producing these files, facilitating a comparative analysis. As for runtime performance, we execute each of the three algorithms on the same computational resource (consisting of 1 node with 10 cores) for genotyping the same sample. We subsequently statistically compared the time taken by each algorithm to genotype individual SNP loci across the sample.

## Declarations

### Availability of data and materials

The original data and software code of this study are available through reasonable request from the corresponding author.

### Author contributions

Yi Wang developed the algorithm and Write the software. Shixuan Zhang and Kanglong Xiao analyzed the data, wrote the main manuscript text. Xinyi He, Meng Hao, Yanyun Ma, Yuxin Tian and Li Jin gave advices in the project. Yi Li and Jiucun Wang devised the project.

## Acknowledgements

We would like to thank the workers, researchers, and participants involved in this study.

## Authors’ information

Shixuan Zhang, Kanglong Xiao have contributed equally to this study.

