## Supplemental information for "A Novel Efficient Algorithm for Common Variants Genotyping from Low-Coverage Sequencing Data"

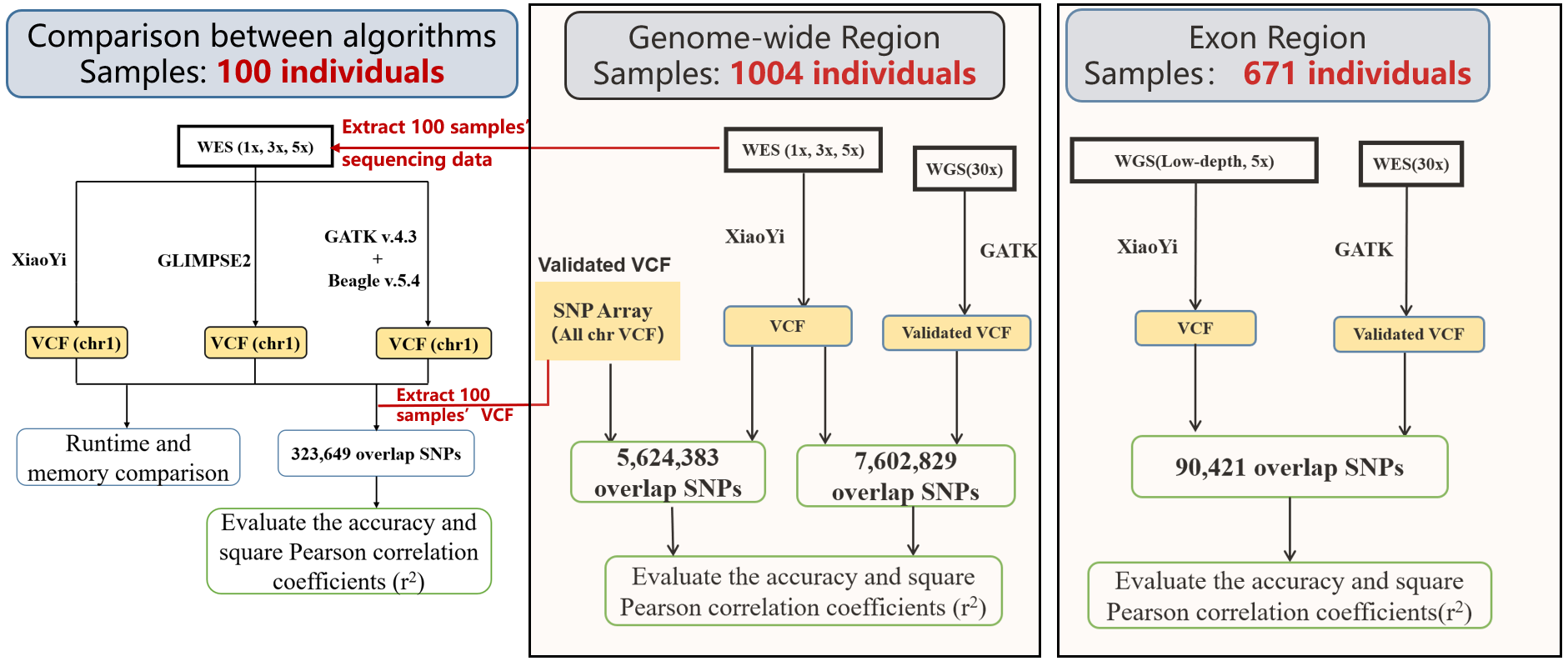

**Fig.S1** **Research Design Flowchart**

**Table S1 Comparison of genotyping performance of three algorithms**

| **MAF Bin (%)** | **accuracy** | | | | | | | | | **r^2^** | | | | | | | | |
| --- | --- | --- | --- | --- | --- | --- | --- | --- | --- | --- | --- | --- | --- | --- | --- | --- | --- | --- |
|  | **GLIMPSE2** | | | **Limpute** | | | **Beagle v.5.4** | | | **GLIMPSE2** | | | **Limpute** | | | **Beagle v.5.4** | | |
|  | **1x** | **3x** | **5x** | **1x** | **3x** | **5x** | **1x** | **3x** | **5x** | **1x** | **3x** | **5x** | **1x** | **3x** | **5x** | **1x** | **3x** | **5x** |
| **(0.02,0.05]** | **0.973** | **0.971** | **0.968** | **0.969** | **0.964** | **0.963** | **0.973** | **0.972** | **0.964** | **0.073** | **0.049** | **0.021** | **0.009** | **0.007** | **0.008** | **0.014** | **0.043** | **0.037** |
| **(0.05,0.1]** | **0.976** | **0.977** | **0.978** | **0.973** | **0.975** | **0.975** | **0.97** | **0.971** | **0.974** | **0.312** | **0.330** | **0.344** | **0.167** | **0.237** | **0.275** | **0.015** | **0.170** | **0.262** |
| **(0.1,0.2]** | **0.979** | **0.98** | **0.98** | **0.976** | **0.979** | **0.979** | **0.968** | **0.971** | **0.974** | **0.463** | **0.479** | **0.485** | **0.308** | **0.406** | **0.436** | **0.016** | **0.276** | **0.363** |
| **(0.2,0.5]** | **0.975** | **0.976** | **0.976** | **0.971** | **0.974** | **0.975** | **0.963** | **0.965** | **0.969** | **0.412** | **0.443** | **0.451** | **0.267** | **0.366** | **0.403** | **0.018** | **0.225** | **0.328** |
| **(0.5,10]** | **0.967** | **0.969** | **0.97** | **0.961** | **0.966** | **0.968** | **0.943** | **0.951** | **0.959** | **0.517** | **0.544** | **0.552** | **0.391** | **0.484** | **0.514** | **0.029** | **0.313** | **0.423** |
| **(1,2]** | **0.962** | **0.965** | **0.966** | **0.958** | **0.964** | **0.965** | **0.923** | **0.942** | **0.952** | **0.603** | **0.625** | **0.630** | **0.538** | **0.607** | **0.625** | **0.042** | **0.414** | **0.516** |
| **(2,5]** | **0.973** | **0.977** | **0.977** | **0.971** | **0.976** | **0.978** | **0.906** | **0.946** | **0.963** | **0.799** | **0.816** | **0.819** | **0.768** | **0.812** | **0.821** | **0.060** | **0.624** | **0.721** |
| **(5,10]** | **0.976** | **0.981** | **0.982** | **0.974** | **0.981** | **0.983** | **0.867** | **0.935** | **0.962** | **0.881** | **0.899** | **0.903** | **0.869** | **0.903** | **0.909** | **0.095** | **0.719** | **0.817** |
| **(10,15]** | **0.972** | **0.979** | **0.98** | **0.972** | **0.98** | **0.982** | **0.823** | **0.919** | **0.954** | **0.906** | **0.923** | **0.927** | **0.901** | **0.929** | **0.934** | **0.140** | **0.752** | **0.843** |
| **(15,20]** | **0.969** | **0.976** | **0.978** | **0.968** | **0.978** | **0.979** | **0.787** | **0.904** | **0.946** | **0.914** | **0.930** | **0.933** | **0.909** | **0.935** | **0.939** | **0.182** | **0.764** | **0.851** |
| **(20,25]** | **0.966** | **0.974** | **0.976** | **0.967** | **0.977** | **0.979** | **0.757** | **0.893** | **0.941** | **0.918** | **0.933** | **0.936** | **0.917** | **0.941** | **0.945** | **0.217** | **0.769** | **0.855** |
| **(25,30]** | **0.964** | **0.972** | **0.974** | **0.964** | **0.976** | **0.977** | **0.721** | **0.877** | **0.934** | **0.921** | **0.936** | **0.938** | **0.919** | **0.944** | **0.947** | **0.259** | **0.763** | **0.852** |
| **(30,35]** | **0.962** | **0.972** | **0.974** | **0.962** | **0.974** | **0.976** | **0.688** | **0.864** | **0.927** | **0.923** | **0.938** | **0.941** | **0.920** | **0.945** | **0.948** | **0.293** | **0.750** | **0.846** |
| **(35,40]** | **0.959** | **0.97** | **0.972** | **0.959** | **0.973** | **0.975** | **0.657** | **0.85** | **0.919** | **0.921** | **0.937** | **0.940** | **0.919** | **0.945** | **0.949** | **0.323** | **0.730** | **0.835** |
| **(40,45]** | **0.961** | **0.971** | **0.973** | **0.96** | **0.973** | **0.976** | **0.62** | **0.841** | **0.915** | **0.927** | **0.942** | **0.945** | **0.924** | **0.947** | **0.951** | **0.353** | **0.725** | **0.833** |
| **(45,50]** | **0.96** | **0.97** | **0.972** | **0.957** | **0.971** | **0.973** | **0.6** | **0.83** | **0.91** | **0.925** | **0.940** | **0.943** | **0.920** | **0.944** | **0.947** | **0.373** | **0.708** | **0.823** |

Comparison of genotyping accuracy and square Pearson correlation coefficients (r^2^) of GLIMPSE2, Beagle v.5.4, and Limpute algorithms with SNPs in different MAF interval segments using microarray data as a validation set.

**Notes:** MAF Bin indicates MAF interval segments, assessed SNP loci are divided according to the MAF values from the public database dbSNPs. 1x,3x,5x represents different sequencing depth.

**Table S2 Genotyping performance measurement of SNPs loci in whole exon region**

| **MAF Bin (%)** | **accuracy** | **r^2^** |
| --- | --- | --- |
| **(0.02,0.05]** | **0.99** | **0** |
| **(0.05,0.1]** | **0.964** | **0.036** |
| **(0.1,0.2]** | **0.905** | **0.232** |
| **(0.2,0.5]** | **0.936** | **0.584** |
| **(0.5,10]** | **0.949** | **0.536** |
| **(1,2]** | **0.945** | **0.617** |
| **(2,5]** | **0.955** | **0.769** |
| **(5,10]** | **0.96** | **0.852** |
| **(10,15]** | **0.957** | **0.869** |
| **(15,20]** | **0.95** | **0.871** |
| **(20,25]** | **0.946** | **0.875** |
| **(25,30]** | **0.943** | **0.869** |
| **(30,35]** | **0.939** | **0.854** |
| **(35,40]** | **0.933** | **0.847** |
| **(40,45]** | **0.932** | **0.834** |
| **(45,50]** | **0.933** | **0.832** |

Genotype accuracy and squared Pearson correlation coefficients (r^2^) of the Limpute algorithm for locus imputation of SNPs in different MAF interval segments using WES data as a validation set.

**Notes:** MAF Bin indicates MAF interval segments, assessed SNP loci are divided according to the MAF values from the public database dbSNPs.

**Table S3 Genotyping performance measurement of SNPs within whole genome regions (based on WGS data as a validation set)**

| **MAF Bin (%)** | **accuracy** | | | **r^2^** | | |
| --- | --- | --- | --- | --- | --- | --- |
|  | **1x** | **3x** | **5x** | **1x** | **3x** | **5x** |
| **(0,0.02]** | **0.971** | **0.974** | **0.976** | **0.185** | **0.210** | **0.230** |
| **(0.02,0.05]** | **0.987** | **0.992** | **0.994** | **0.314** | **0.503** | **0.613** |
| **(0.05,0.1]** | **0.984** | **0.99** | **0.993** | **0.414** | **0.609** | **0.705** |
| **(0.1,0.2]** | **0.978** | **0.988** | **0.991** | **0.555** | **0.736** | **0.811** |
| **(0.2,0.5]** | **0.967** | **0.983** | **0.989** | **0.710** | **0.849** | **0.895** |
| **(0.5,10]** | **0.957** | **0.98** | **0.986** | **0.815** | **0.913** | **0.940** |
| **(1,2]** | **0.957** | **0.98** | **0.986** | **0.876** | **0.943** | **0.960** |
| **(2,5]** | **0.966** | **0.984** | **0.988** | **0.905** | **0.954** | **0.966** |
| **(5,10]** | **0.967** | **0.984** | **0.988** | **0.911** | **0.955** | **0.966** |
| **(10,15]** | **0.963** | **0.982** | **0.986** | **0.915** | **0.956** | **0.965** |
| **(15,20]** | **0.958** | **0.979** | **0.984** | **0.917** | **0.955** | **0.963** |
| **(20,25]** | **0.954** | **0.976** | **0.981** | **0.914** | **0.950** | **0.958** |
| **(25,30]** | **0.95** | **0.974** | **0.979** | **0.910** | **0.946** | **0.954** |
| **(30,35]** | **0.948** | **0.972** | **0.978** | **0.904** | **0.941** | **0.948** |
| **(35,40]** | **0.947** | **0.97** | **0.976** | **0.898** | **0.934** | **0.941** |
| **(40,45]** | **0.945** | **0.968** | **0.974** | **0.890** | **0.927** | **0.934** |
| **(45,50]** | **0.944** | **0.968** | **0.974** | **0.886** | **0.924** | **0.931** |

Genotype accuracy and squared Pearson correlation coefficients (r^2^) of the Limpute algorithm for imputation of SNPs loci in different MAF interval segments using WGS data as a validation set.

**Notes:** MAF Bin indicates MAF interval segments, assessed SNP loci are divided according to the MAF values from the public database dbSNPs. 1x,3x,5x represents different sequencing depth.

**Table S4 Genotyping performance measurement of SNPs loci in whole genome regions (based on microarray data as validation set)**

| **MAF Bin (%)** | **accuracy** | | | **r^2^** | | |
| --- | --- | --- | --- | --- | --- | --- |
|  | **1x** | **3x** | **5x** | **1x** | **3x** | **5x** |
| **(0.02,0.05]** | **0.963** | **0.968** | **0.969** | **0.006** | **0.042** | **0.058** |
| **(0.05,0.1]** | **0.965** | **0.973** | **0.974** | **0.049** | **0.189** | **0.234** |
| **(0.1,0.2]** | **0.964** | **0.973** | **0.976** | **0.082** | **0.276** | **0.330** |
| **(0.2,0.5]** | **0.957** | **0.969** | **0.972** | **0.114** | **0.314** | **0.370** |
| **(0.5,10]** | **0.942** | **0.961** | **0.964** | **0.210** | **0.417** | **0.465** |
| **(1,2]** | **0.936** | **0.956** | **0.96** | **0.360** | **0.537** | **0.573** |
| **(2,5]** | **0.95** | **0.967** | **0.971** | **0.604** | **0.728** | **0.755** |
| **(5,10]** | **0.956** | **0.972** | **0.975** | **0.747** | **0.833** | **0.851** |
| **(10,15]** | **0.952** | **0.969** | **0.972** | **0.802** | **0.866** | **0.879** |
| **(15,20]** | **0.948** | **0.965** | **0.969** | **0.829** | **0.881** | **0.891** |
| **(20,25]** | **0.943** | **0.962** | **0.965** | **0.843** | **0.887** | **0.896** |
| **(25,30]** | **0.939** | **0.958** | **0.962** | **0.848** | **0.888** | **0.895** |
| **(30,35]** | **0.937** | **0.956** | **0.96** | **0.854** | **0.891** | **0.898** |
| **(35,40]** | **0.935** | **0.953** | **0.957** | **0.853** | **0.887** | **0.893** |
| **(40,45]** | **0.933** | **0.952** | **0.956** | **0.856** | **0.889** | **0.894** |
| **(45,50]** | **0.933** | **0.951** | **0.956** | **0.855** | **0.886** | **0.892** |

Genotype accuracy and squared Pearson correlation coefficients (r^2^) of the Limpute algorithm for imputation of SNPs loci in different MAF interval segments using microarray data as a validation set.

**Notes:** MAF Bin indicates MAF interval segments, assessed SNP loci are divided according to the MAF values from the public database dbSNPs. 1x,3x,5x represents different sequencing depth.
